# CRISPR/Cas9-Mediated Cldn2 Knockout in HCT116 Cells, Reveals Its Crucial Role in Colorectal Cancer progression

**DOI:** 10.1101/2025.02.28.640859

**Authors:** Rana A. Alghamdi, Maryam H. Al-Zahrani

## Abstract

**Background:** Colon cancer progression heavily relies on intricate mechanisms of invasion, metastasis, and migration. Tight junction protein Cldn2 has emerged as a potential regulator of these processes. This study aimed to elucidate the molecular mechanisms linking Clan2 deletion to gene expression changes related to motility, invasion, and metastasis in colon caner.

**Methods:** CRISPR/Cas9-mediated knockout of human Cldn2 in HCT116 cells was conducted, and the resulting cells were compared to the wild-type cells using real-time PCR to analyze the expression of genes associated with invasion and metastasis.

**Results:** Cldn2-KO resulted in a widespread downregulation of genes linked to motility, invasion, and metastasis, including ZONAB, NDRG1, Cldn14, Cldn23, Bcl2, , P53, and BCL-6. These findings suggest a potential regulatory role of Cldn2 in the expression of these genes, influencing colon cancer cell migration and spread.

**Conclusion:** This study identified Claudin-2 as a crucial regulator of genes involved in colorectal cancer metastasis. Downregulation of these genes upon Claudin-2 deletion suggests its inhibitory role in cancer cell motility and invasion. Further investigation into the specific downstream signaling pathways mediated by Claudin-2 could pave the way for novel therapeutic strategies targeting metastasis inhibition.

## 1. Introduction

Colorectal cancer (CRC) is a deadly disease influenced by both environmental and genetic factors. It considered one of the most prevalent malignant tumors is, which is becoming more widespread in eastern regions (Siegel et al., 2023). Proteomic and genomic methods have changed the field of cancer research recently and have given important new understandings of the molecular pathways behind colorectal cancer (CRC) (Sarkar, 2023). It has been revealed that Claudin-2 (Cldn2), a constituentof the cellular tight junction, contributes to the advancement of different types of malignancyby exhibiting aberrant expression (Tabariès et al., 2012).

Claudin family plays a key role in cell barriers, differentiation, and proliferation. The expression patterns of Claudin knockout varied throughout malignancies and organs (Cherradi *et al*., 2017). Thus, Cldns have been proposed as targets for cancer treatment as well as diagnostic indicators. A previous work, has shown that there is increasing agreement on the potential value of Claudin-2 as a biomarker of prognostic and therapeutic features in colorectal cancer (Alghamdi and Al-Zahrani, 2023). Interestingly, Claudin-2 (Cldn2) is a part of biological junctions that are tight and was first shown to be abundant in inflammatory bowel disease (IBD) and or colorectal cancer (CRC) tissues (Weber *et al*., 2008). Cldn2 is the most distinct member of the family and is exhibits a unique expression pattern because its expression is limited to permeable epithelial tissues (Rosenthal *et al*., 2010).

In colon cancer, Cldn2 appears to be an oncogene that affects its growth and migration/invasion via EGFR-mediated signaling transactivation (Dhawan *et al*., 2011). The increasing incidence of CRC has been connected to the presence of Cldn2 in cellular tight junctions. It has been demonstrated that Cldn2 inhibits NDRG1 transcription, thereby promoting CRC development and metastasis (Wei *et al*., 2021). When comparing neighboring normal tissues to most CRC tissues, high expression of Cldn2 has been found, and this has been linked to decreased cancer-specific survival rates. In addition, individuals with stage II/III CRC receiving adjuvant treatment had a worse prognosis and an increased risk of recurrence when their expression of Cldn2 was high (Al-Zahrani *et al*., 2016). Although environmental and genetic variables can affect CRC, knowing how Cldn2 functions in the disease’s course could help identify new treatment options. High Claudin-2 expression patterns showed shorter survival outcomes indicating a connection between high Claudin-2 expression and CRC progression (Wei *et al*., 2021).

While genomic sequencing has shed light on mutations driving colon cancer development, understanding the precise roles of these alterations and their complex interplay with cellular pathways has been a persistent challenge (Kim and Cho, 2023). Gene editing technology, particularly the CRISPR-Cas9 system, has emerged as a promising approach for studying cancer Furthermore, CRISPR-Cas9 enables the creation of genetically defined colon cancer models, replicating the diverse landscape of the disease (Zhao et al., 2021).

The use of gene expression profiling and CRISPR-Cas9 technology has allowed researchers to gain a deeper and more comprehensive understanding of colon cancer, its pathways, and potential therapeutic targets. By combining gene editing techniques with the analysis of biomarker data, scientists can detect novel mutations in CRC that confer resistance or sensitivity to drugs, paving the way for the development of more effective treatments (Meng et al., 2023). Hence, the utilization of biomarker signatures may have a crucial or conclusive impact on the advancement of tailored medical healthcare. Given the limited understanding of Cldn2’s regulatory mechanisms and its role in CRC progression, we employed CRISPR-Cas9 editing to dissect its impact on gene expression and metastasis potential.

## 2. Materials and Methods

### 2.1. Cell culture

Wild type and knockout HCT116 cells were cultured in McCoy’s 5A medium (16600082, Thermo Scientific, MA, USA) supplemented with 10% Fetal Bovine Serum (10270106, Thermo Scientific, MA, USA) and 1% Penicillin-Streptomycin (15140-122, Thermo Scientific, MA, USA). The cells were maintained at 37°C in a humidified incubator with 5% CO2.

### 2.2. Generation of CRISPR/Cas9-mediated Cldn2 knockout (KO) cells

CRISPR Cas9 mediated knockout of human Cldn2 (gene ID: 9075) in (HCT116) cells were produced by Synthego Corporation (Redwood City, CA, USA).

The generation of these cells involves the use of Ribonucleoproteins that consist of the Cas9 protein and synthetic chemically modified single-guide RNA (GGUGCUAUAGAUGUCACACU) against exon 2 of human Cldn2 (gene ID: 9075) were electroporated into the cells. Guide RNA cut location: chrX:106,928,419, followed by single-cell FACS sorting and validation (Enzmann and Wronski, 2018). The assessment of editing efficiency takes place 48 hours after electroporation. The process involves extracting genomic DNA from a subset of cells, followed by PCR amplification and sequencing using the Sanger sequencing method. PCR Primers: F:CAGCCTGAAGACAAGGGAGC, R: TGTCTTTGGCTCGGGATTCC. Sequencing Primer Used: CAGCCTGAAGACAAGGGAGC

The chromatograms obtained are analyzed using the Synthego Inference of CRISPR edits software (ICE) (Roginsky, 2018).

For insertion-deletion The indel examination was conducted by Synthego using Illumina sequencing. A total of 20 ng of genomic DNA (gDNA) was utilized as a template for amplifying the region surrounding the sgRNA target site, using the specified primers.

In order to generate monoclonal cell populations, cell pools with altered Cldn2 gene (Cldn2-KO) are distributed at a density of 1 cell/well using a single cell printer onto either 96 or 384 well plates. Every 3 days, all wells are recorded to verify the growth of a clone originating from a single cell. The PCR-Sanger-ICE genotyping technique is used to screen and identify clonal populations.

### 2.3. Wound healing assay

The migration capacity of wild-type (Wt) and Claudin-2 knockout (Cldn2-KO) cells was assessed using a wound healing assay. Briefly, 30×10^4^ cells per well were seeded in 6-well plates and cultured until reaching 90% confluence. A sterile pipette tip was then used to scratch across the center of each well, creating a defined wound area. Detached cells were removed by 500 μL PBS wash, then fresh culture medium was added. Subsequently, cell migration into the scratched area was monitored and captured at 0, and 24 hours using a Nikon E 600 phase-contrast microscope. Images of the wound area were analyzed using ImageJ software. Wound closure percentage was calculated as 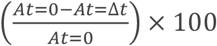, where A_t=0_ is scratch area in 0 time, At=_Δt_ is scratch area in 24h.

### 2.4. RNA extraction and quantitative RT-PCR

RNA was isolated in accordance with the manufacturer’s protocol using an RNA extraction kit (Cat# R1200, Suolaibao, China). Subsequently, cDNA was synthesized using random nonamer primers and the First-Strand Synthesis System (Sigma-Aldrich, United States). EvaGreen fluorescence-based real-time PCR was performed using reagents purchased from Applied Biological Materials (Richmond, Canada). The RT-qPCR primers used in this study are listed in Table 1. The gene expression levels in the samples were investigated using RT-qPCR, and the cDNA synthesis and PCR procedures were optimized to ensure high-quality results.

**Table 1:**
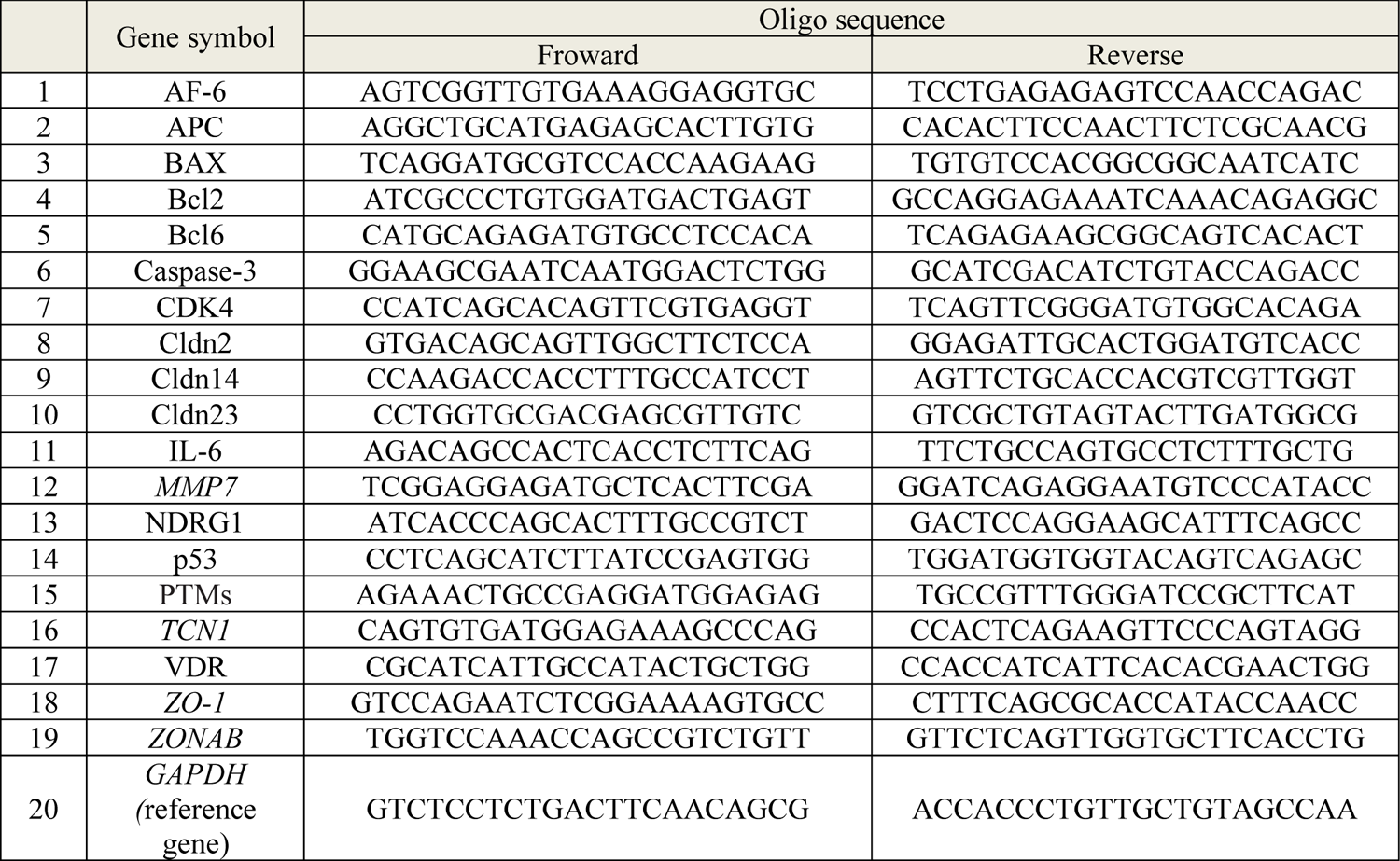
Primers for RT-qPCR used in this study. RG: reference gene.

### 2.5. Relative and normalized fold expression values calculation

The gene expression patterns of several target genes were examined in samples from both wild-type (Wt) and knockout (Cldn2-KO) individuals using real-time PCR. For every target gene, the Ct values that were derived from the amplification curves were used to compute ΔCt, copy number, ΔΔCt, and fold change.

ΔCT was calculated using formula:

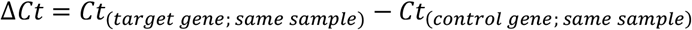

Copy number was calculated using formula:

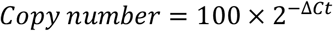

ΔΔCt was calculated using formula:

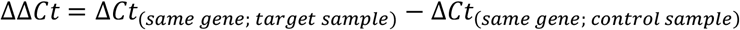

Fold change was calculated using formula:

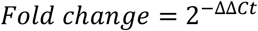

The housekeeping gene GAPDH was used as an internal control for normalization. The expression levels of the housekeeping gene GAPDH were relatively stable across all samples

### 2.6. Statistical analysis

Statistical analyses were conducted using unpaired t-tests for normally distributed data and Mann-Whitney tests for non-normal data. Data are presented as mean ± SEM, all experiments were repeated at least three times, and analyses were performed using GraphPad Prism.

## 3. Results

### 3.1. High efficiency of CRISPR editing Cldn2 gene

The Synthego ICE CRISPR Analysis tool was employed to assess the effectiveness of CRISPR Cldn2 gene knockout as shown in figure 1. The ICE value of 91% indicates a high degree of CRISPR editing efficiency, suggesting that a substantial proportion of the cells within the edited population harbored the intended Cldn2 gene knockout with 91% of insertion-deletion (INDL) of Cldn2 gene after CRISPR editing.

**Figure 1.**
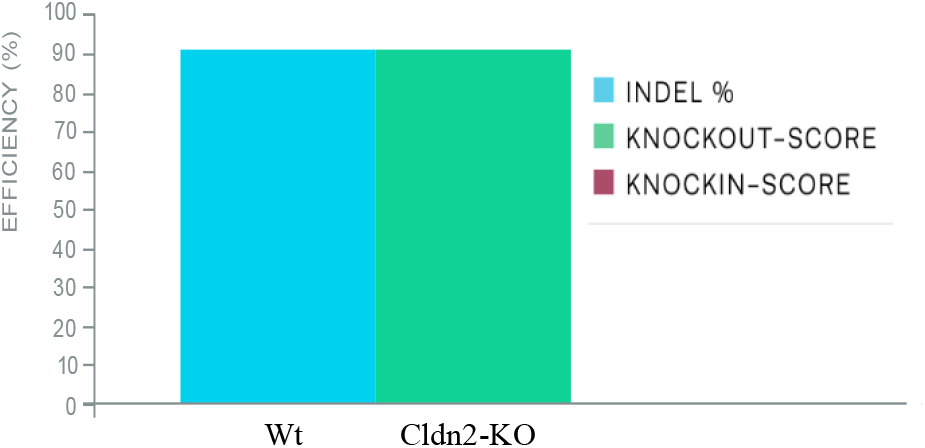
CRISPR editing efficiency of Cldn2-KO, This bar graph shows that 91% of the cells with contain the desired mutation with 91 % of INDLof Cldn2 gene after CRISPR editing.

The discordance plot in figure 2, provides a visual representation of the alignment between Wt and Cldn2-KO within the inference window, which encompasses the region surrounding the CRISPR cut site. It highlights the degree of sequence discordance, indicating the average level of deviation from the reference sequence.

**Figure 2.**
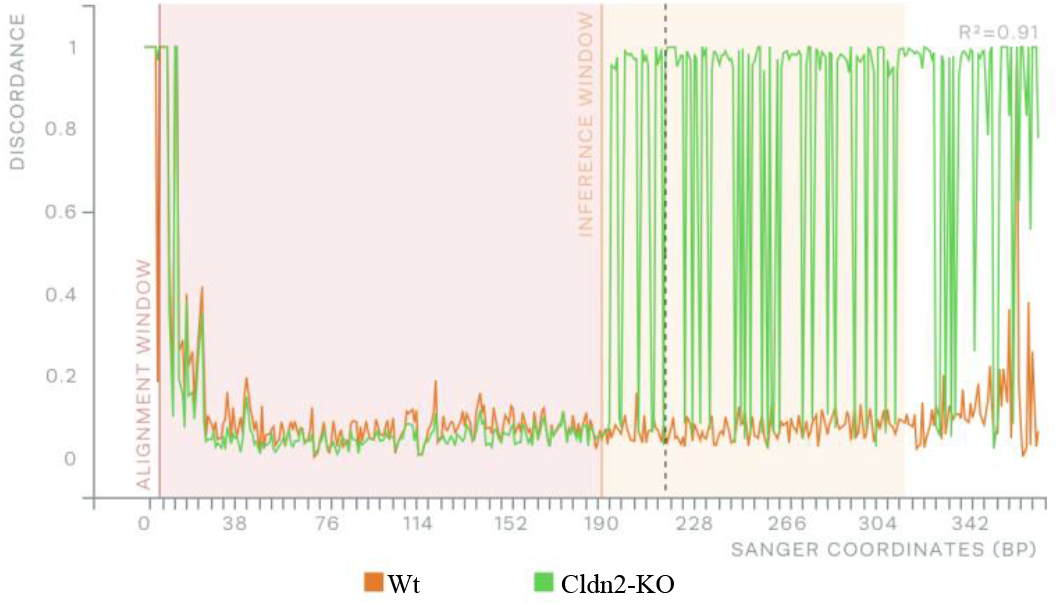
Sanger discordance plot shows the alignment per base between the Wt and Cldn2-KO samples, inside the inference window (the area around the cute site), indicates the average proportion of signal that differs from the reference sequence obtained from the Wt trace. In the plot, the proximity between the green line and orange line is seen before to the cut site. However, a typical CRISPR edit leads to a significant increase in sequence discordance at the cut site, causing the green and orange lines to stay far apart fterwards.

**Figure 3.**
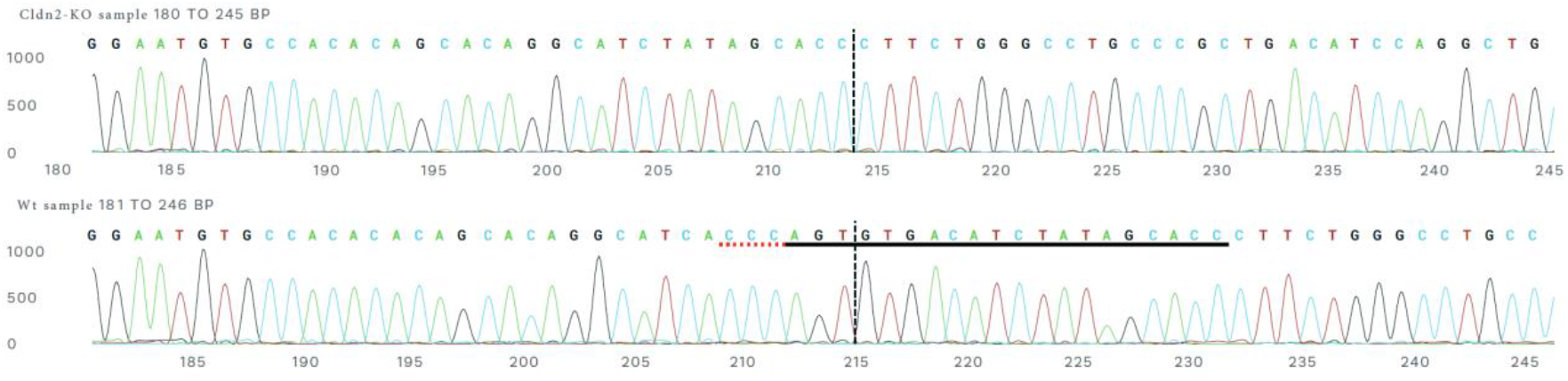
The Sanger sequence view displays the knockout and Wt sequences in the immediate area of the guide sequence. This displays the sequence base calls obtained from both the Wt and the experimental sample. The guide sequence is represented by the horizontal black underlined section. The PAM location is indicated with a horizontal red underline. The vertical line, shown with black dots, depicts the precise location of the incision. Performing a cut and attempting to repair it often leads to the presence of mixed sequencing bases.

**Figure 3a.**
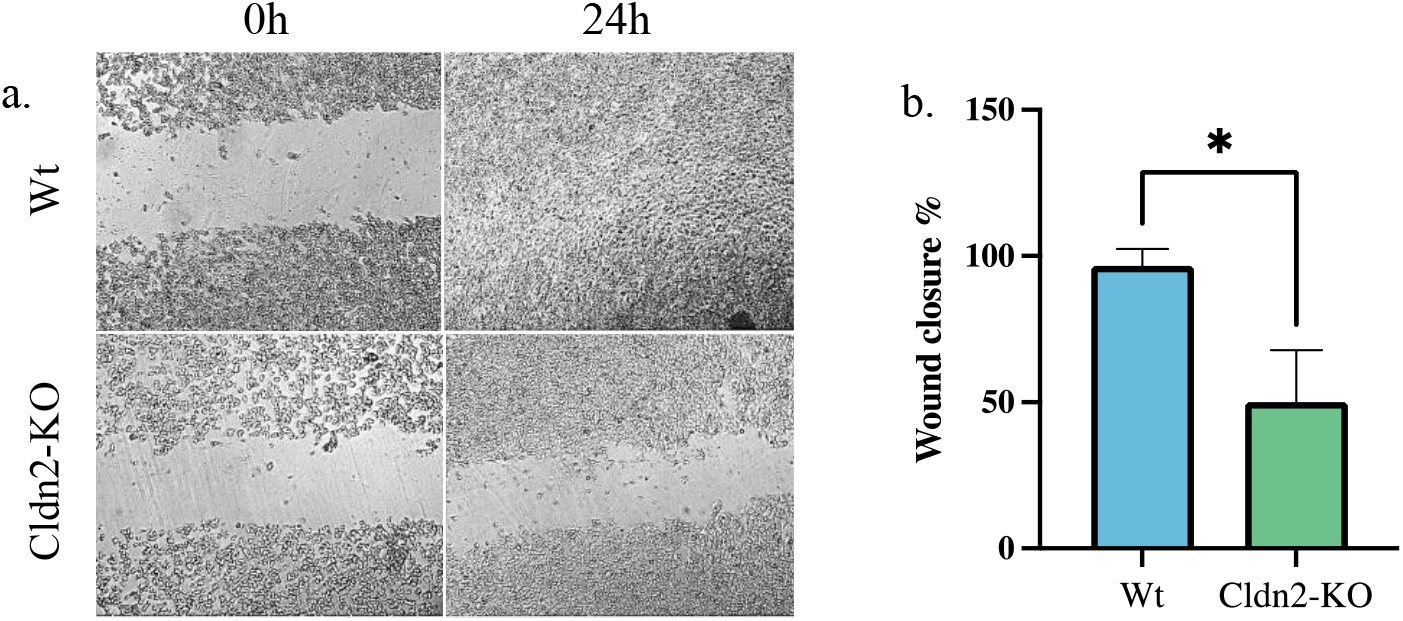
Effect of Cldn2 knockout on HTC116 cell migration: a) Inhibition of HTC116 cell migration in response to Cldn2 knockout over 24h. b. The percentage of wound closure was significantly lower in Cldn2-KO at 41% compared with Wt at 96%. Data show mean ± SEM. \**P* < 0.005.

**Figure 3b.**
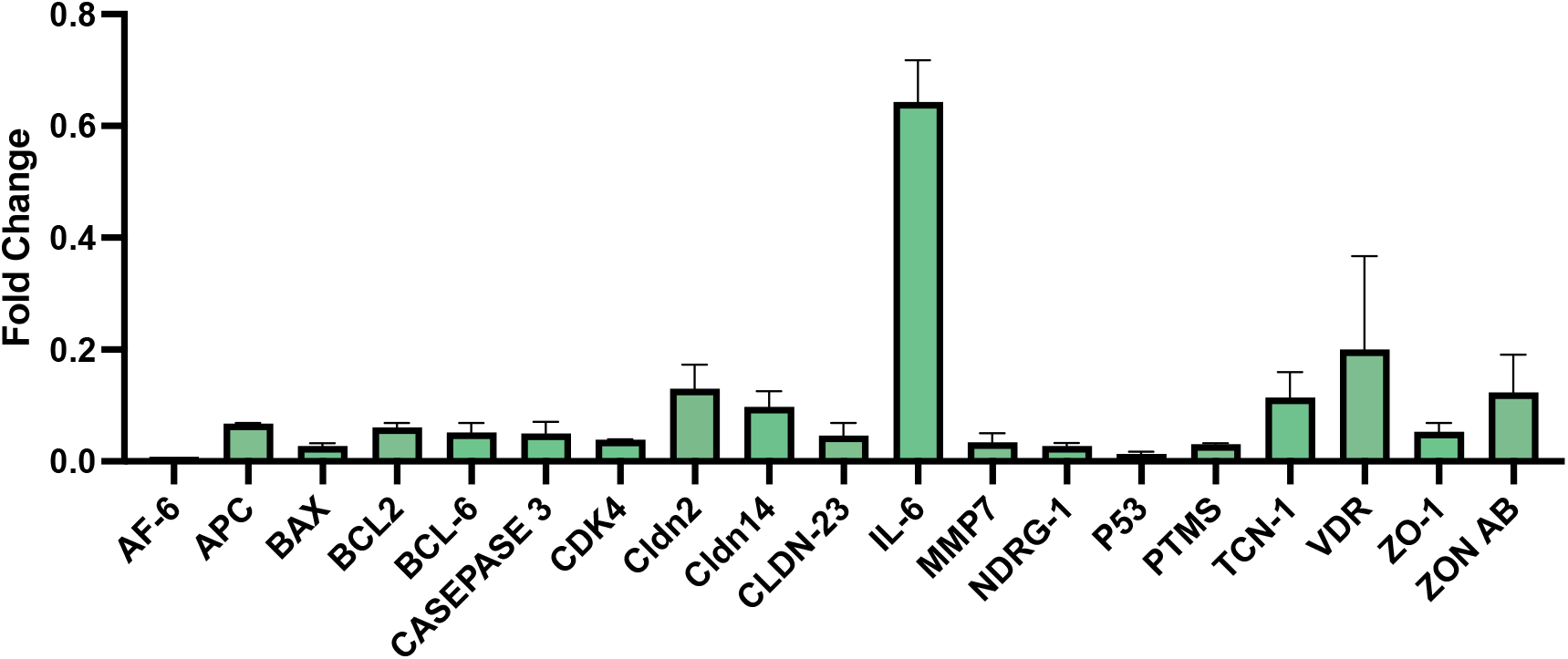
Comparison of Fold Change Across Multiple Genes in Wild-type (Wt) and Knockout (Ko) Samples: The X-axis represents different groups, while the Y-axis illustrates the copy number estimates. IL-6 exhibits the lowest level of downregulation in Cldn2-KO samples, with a fold change of 0.718, AF-6 shows the most substantial decrease, with a fold change of 0.008.

### 3.2. Migration capacity of HCT16 cells was abolished by Cldn2 knockout

The effect of Cldn2 knockout on HTC116 cell migration was shown in Fig 4a. The wound closure rate of HCT116 cell in response to Cldn2 knockout was notablydecreased over 24h. The percentage of wound closure (Fig 4b) was significantly lower in Cldn2-KO at 41% compared with Wt at 96%. Cldn2-KO cells were significantly inhibited by (p value = 0.0027).

**Figure 4.**
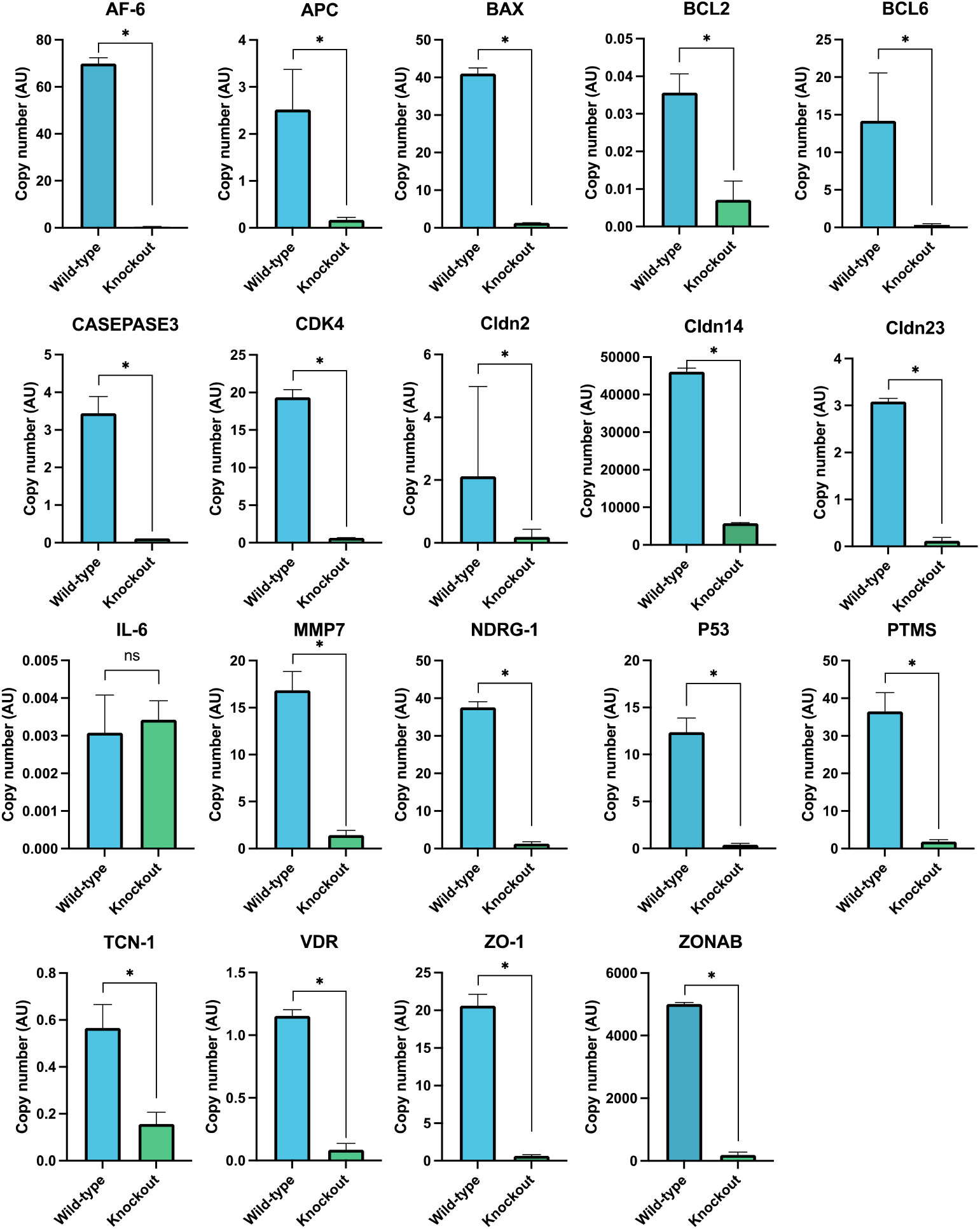
Gene expression profiles in Wt and Cldn2-KO samples, each bar in the chart corresponds to a specific gene, and the height of the bars indicates the magnitude of fold change observed in the Cldn2-KO samples compared to the Wt samples. Data show mean ± SEM. *P < 0.005, ns= not significant.

### 3.3. Gene expression profiles in Wt and Cldn2-KO samples

We profiled the gene expression in Wt and Cldn2-KO cells using results from three independent experiments. Figure 4 shows the copy number variations of target genes in Wt and Cldn2-KO cells. Analysis of expression patterns for all genes consistently showed a considerable decrease in expression in Cldn2-KO samples compared to the Wt. The observed patterns indicates that the knockout of Cldn2 has a significant impact on its interacting partners causing their downregulation.

Figure 5 depicts the fold change values of several genes in both Wt and Cldn2-KO samples. Each bar in the chart corresponds to a specific gene, and the height of the bars indicates the magnitude of fold change observed in the Ko samples compared to the Wt samples. The highest bars on the figure represent the gene with the lowest degree of expression variation. Of all the genes that were evaluated, IL-6 is the least downregulated in Cldn2-KO samples, with a fold change of 0.718 and maintains a relatively stable expression pattern. On the other hand, AF-6 AFADIN exhibits the most significant downregulation, with a fold change of 0.008.

## 4. Discussion

Claudins are prominent transmembrane proteins that are present in tight junctions of both epithelial and endothelial tissues ( Venugopal, Anwer and Szászi, 2019). Studies suggested that claudins involved in the development of malignancies, however the specific process is not yet fully understood (Zhu et al., 2019; Al-Zahrani et al., 2023; Alghamdi and Al-Zahrani, 2023). The significance of Cldn2 in tumor development was shown by its significant contribution to the motility, invasion, and metastasis of cancer cells. Despite extensive investigation in both experimental and clinical settings, the specific role and underlying mechanism of Cldn2 in CRC are still not well understood (Rosenthal et al., 2010; Dhawan et al., 2011; Tabariès et al., 2021; Wei et al., 2021; Ahmad et al., 2023).

The progression of colon cancer is largely dependent on the complex mechanisms of invasion, metastasis, and migration. Tight junction proteins, especially Cldn2, have come to light as possible regulators of these processes among the many molecular mediators. In order to better understand the molecular mechanisms causing colon cancer metastasis, current study investigated how Cldn2 deletion affects the expression of genes linked to motility, invasion, and metastasis in HCT 116 cells. Examining all genes associated with invasion and metastasis revealed a regular pattern of downregulation in response to Cldn2 knockdown in HCT 116 cells.

Previous research indicates that suppression of Cldn2 expression leads to the disassembly of the Cldn2/ZO1/ZONAB complex, which results in ZONAB translocating to the nucleus and promoting NDRG1 transcription (Tao *et al*., 2023). Significant ZONAB enrichment was seen in HT29 cells at the NDRG1 promoter region in Cldn2 downregulated cells, but a notable reduction was observed in Cldn2 overexpressed HCT116 cells. On the other hand, our research indicates that both ZONAB and NDRG1 are downregulated when Cldn2 is not present. ZO-1 has a crucial role in controlling the activity of Cldn2. Previous findings suggest that the abundance of Cldn2 is regulated by confluence (mature junctions) and the ZO-1 protein at various degrees. The existence of fully developed tight junctions (TJs) and ZO-1 protein decreases the degradation of Cldn2. Furthermore, they enhance the activity of the Cldn2 promoter (Amoozadeh *et al*., 2018). In the present study, the downregulation of ZO-1 in Cldn2 in knockout cells is a marker that downstream signaling is disrupted in the absence of Cldn2.

Both Cldn14 and Cldn23 have been implicated in promoting the growth and migration of cancer cells. Cldn14 has been directly associated with the stimulation of the PI3K/AKT/mTOR pathway, which plays a role in promoting the proliferation and migration of cancer cells (Wei, Xiao-nan and Jin, 2018; Tian-Yu *et al*., 2021). In one research, it was shown that at the mRNA level, the expression of Cldn2 and Cldn14 was increased, whereas the expression of Cldn23 was decreased in patients with colorectal cancer, CRC (Yang *et al*., 2022). In the current study, both Cldn14 and Cldn23 show downregulation in the knockout condition, suggesting a potential regulatory role of Cldn2 in their expression.

In spontaneous CRC, Cldn2 is found at the base of the crypt, forms clusters with Cyclin-D1 and c-Myc proteins, and controls the proliferation, differentiation, and migration of intestinal epithelial cells (IECs) in along with cancer stem cells. Notably, Cldn2 expression is increased in CRC and facilitates its progression (Mariadason *et al*., 2005; Dhawan *et al*., 2011; Ahmad *et al*., 2017; Paquet-Fifield *et al*., 2018; Tabariès *et al*., 2021). Previous studies report a notable decrease in the expressions of Bcl2 and C-Myc in Cldn2-KO mice, which leads to a compromised ability to recover from colitis and an increased susceptibility to intestinal damage caused by various ways (Ahmad *et al*., 2023). Our results are in accordance with these findings as Bcl2 is found to be downregulated in Cldn2-KO cell lines.

The inverse relationship between p53 and Cldn2 has been documented in mice colon tissues induced with dextran sulfate sodium colitis (Kim *et al*., 2020). Furthermore, a study proposed a negative regulatory complex involving p53 and HNF4α that may impact Cldn2 expression (Hirota *et al*., 2021). However, in the present study the downregulation of P53 in the knockout condition, as indicated by the fold change value of 0.02, suggests that Cldn2 may play a regulatory role in its expression. Reduced copy number provides additional evidence in favor of the notion of downregulation in the Cldn2 knockout.

A recent study provides an evidence for the qualitative and quantitative expression of Bcl6 involvement in human CRC progression, and demonstrate that *Bcl6* appears to be involved in tumor development, from the earliest stage of carcinogenesis (Sena *et al*., 2014). In our study, in the knockout condition, Bcl6 has shown a downregulation, pointing to a possible regulatory function for Cldn2 in its expression. Cellular signaling cascades may be affected by Cldn2 inhibition, which may compromise tight junction integrity. Downstream genes, including BCL6, may be impacted by this disturbance in terms of expression.

AF-6, also known as Afadin, is a pivotal adhesion protein that orchestrates tight junction (TJ) formation and cell-cell adhesion (Niessen, 2007). Studies have shown that CLDN2 promotes AF-6 recruitment to TJs, while AF-6, in turn, stabilizes CLDN2 at the cell membrane, this reciprocal interaction reinforces TJ integrity and cell adhesion (Wang *et al*., 2022). Interestingly, our results demonstrate that Cldn2 knockout coincides with a downregulation of AF-6 expression. This observation aligns with previous reports highlighting a co-dependent relationship between AF-6 and Cldn2, as its found to be Claudin-2 interacting partner that help breast cancer cells proliferate in soft agar and spread to the liver and/or lungs (Zhang *et al*., 2018; Tabariès *et al*., 2019). Breast and colorectal cancer (CRC) liver metastases may be related to similar signaling pathways downstream of Claudin-2-Afadin (Tabariès *et al*., 2021). Low afadin expression was linked to lower levels of claudin-2 and a higher risk of pulmonary metastases in individuals with osteosarcoma (Zhang *et al*., 2018). Downregulatory patterns were observed for Afadin as well in the present research.

The IL-16 cytokine is encoded as a precursor molecule known as pro-IL-16. This molecule undergoes cleavage by caspase-3, resulting in the production of a smaller secreted molecule. The levels of IL-16 showed a direct correlation with the progression of gastrointestinal tumors. Additional research has shown that there are elevated levels of IL-16 in both malignant ovarian tumors and cutaneous T-cell lymphoma, as evidenced by increased tissue expression and serum concentrations (Blaschke *et al*., 1999; Asadullah *et al*., 2000; Yellapa *et al*., 2014). In AOM/DSS-treated Cldn2KO-mice, there were significant increases in the expression of caspase-3, a hallmark of DNA damage and apoptosis (Ahmad *et al*., 2023). However, Low expression levels were observed for IL-16 and caspase-3 in the present study.

TCN1 has high expression levels in tumor tissues, and this expression is positively associated with the severity of cancer. TCN1 may be a key oncogene for colorectal cancer, according to reports (Li, Guo and Cai, 2021). In one investigation, clinical colorectal carcinoma samples showed elevated expression of TCN1, and there was a positive association between their levels. Through its interaction with ITGB4, it has been shown that the lack of TCN1 in colorectal cancer (CRC) cells inhibits cell proliferation, adhesion, and invasion (Zhu *et al*., 2023). Contrary to this,TCN1 was found to be downregulated in the Cldn2 knock out cells in the present study.

The regulation of claudins by the adenomatous polyposis coli (APC) tumor suppressor protein is intricately linked to the Wnt signaling pathway. APC gene mutations are recognized for stabilizing β-catenin, an essential element in the Wnt pathway, and are crucial in the oncogenic development of intestinal epithelial cells, which may result in the development of carcinomas and adenomas (Miwa *et al*., 2001; Mankertz *et al*., 2004). Activation of Wnt signaling has been shown in around 30%–50% of stomach tumors in humans. The expression of MMP7 is increased at both the mRNA and protein levels, and this is associated with the activation of the Wnt-β-catenin pathway in TNBC (Triple Negative Breast Cancer). Additional patient data reveals that MMP7 is increased in a particular subgroup of TNBC with a distinct feature of reduced PTEN expression, which is a gene responsible for suppressing tumor growth (Dey *et al*., 2013). APC and MMP7 were found to be low in expression in Cldn2 knock outs.

According to one study, VDR inhibits Claudin-2 promoter activity by a mechanism that is reliant on the presence of a Cdx1 binding site. Claudin-2 levels decrease at the mRNA and protein levels when intestinal VDR is lacking in epithelial cells (Zhang *et al*., 2015). The lack of Vitamin D receptor (VDR) in the intestine causes an imbalance in the processes of cell self-destruction (apoptosis) and self-renewal (autophagy). This imbalance is marked by an increase in cell death and higher levels of the proapoptotic protein BAX. However, both BAX and VDR are found to be downregulated in the present study in the absence of Cldn2.

CDK4, or cyclin-dependent kinase 4, is a crucial enzyme involved in regulating cell division and preventing cell death. Its role in colon cancer has been extensively studied, with some studies showing that CDK4 activation leads to increased tumor growth, while others have found that knocking out CDK4 reduces adenoma development (Thoma, Neurath and Waldner, 2021). Research has shown that CDK4 expression levels are associated with the expression of apical junction proteins, including Cldn2, in colon cancer (González-Mariscal *et al*., 2020). Our findings suggest that CDK4 and CLDN2 may be involved in the progression of colon cancer, with potential implications for the development of targeted therapeutic strategies. However, further research is needed to fully elucidate the relationship between CDK4 expression and colon cancer in the context of CLDN2 knockout.

## 5. Conclusions

In conclusion, the present study demonstrates a crucial role for Cldn2 in regulating genes associated with CRC metastasis. Disruption of Cldn2, a key tight junction protein, significantly altered HCT16 cell migration through its effects on critical signaling pathways. These findings uncover an intricate network of interactions governed by Cldn2, suggesting its potential as a valuable target for both diagnostic and therapeutic interventions in colon cancer. However, further investigation is necessary to elucidate the precise mechanisms by which Cldn2 governs cell migration and its potential interactions with other genetic factors in a clinical setting.

## Disclosure statement

No potential conflict of interest was reported by the author(s).

## Notes

### Competing Interest Statement

The authors have declared no competing interest.

## References

Ahmad, R. et al. (2017) ‘HDAC-4 regulates claudin-2 expression in EGFR-ERK1/2 dependent manner to regulate colonic epithelial cell differentiation’, Oncotarget. United States, 8(50), pp. 87718–87736. doi: 10.18632/oncotarget.21190.

Ahmad, R. et al. (2023) ‘Claudin-2 protects against colitis-associated cancer by promoting colitis-associated mucosal healing’, The Journal of clinical investigation. United States, 133(23), p. e170771. doi: 10.1172/JCI170771.

Al-Zahrani, M. et al. (2016) ‘Evaluation of Prognostic Potential of CD44 , Claudin-2 and EpCAM Expression Patterns in Colorectal Carcinoma’, 10(March), pp. 147–156.

Al-Zahrani, M. H. et al. (2023) ‘Expression pattern, prognostic value and potential microRNA silencing of FZD8 in breast cancer’, Oncology letters. Greece, 26(5), p. 477. doi: 10.3892/ol.2023.14065.

Alghamdi, R. A. and Al-Zahrani, M. H. (2023) ‘Identification of key claudin genes associated with survival prognosis and diagnosis in colon cancer through integrated bioinformatic analysis’, Frontiers in Genetics. Frontiers, 14, p. 1221815. doi: 10.3389/fgene.2023.1221815.

Amoozadeh, Y. et al. (2018) ‘Cell confluence regulates claudin-2 expression: possible role for ZO-1 and Rac’, American Journal of Physiology-Cell Physiology. American Physiological Society, 314(3), pp. C366–C378. doi: 10.1152/ajpcell.00234.2017.

Asadullah, K. et al. (2000) ‘IL-15 and IL-16 overexpression in cutaneous T-cell lymphomas: stage-dependent increase in mycosis fungoides progression’, Experimental Dermatology. Wiley, 9(4), pp. 248–251. doi: 10.1034/j.1600-0625.2000.009004248.x.

Blaschke, V. et al. (1999) ‘Expression of the CD4+ Cell-Specific Chemoattractant Interleukin-16 in Mycosis Fungoides’, Journal of Investigative Dermatology. Elsevier BV, 113(4), pp. 658–663. doi: 10.1046/j.1523-1747.1999.00717.x.

Cherradi, S. et al. (2017) ‘Antibody targeting of claudin-1 as a potential colorectal cancer therapy’, Journal of experimental & clinical cancer research : CR. England, 36(1), p. 89. doi: 10.1186/s13046-017-0558-5.

Dey, N. et al. (2013) ‘Differential activation of Wnt-β-catenin pathway in triple negative breast cancer increases MMP7 in a PTEN dependent manner’, PloS one. United States, 8(10), pp. e77425–e77425. doi: 10.1371/journal.pone.0077425.

Dhawan, P. et al. (2011) ‘Claudin-2 expression increases tumorigenicity of colon cancer cells: role of epidermal growth factor receptor activation’, Oncogene. 2011/03/07, 30(29), pp. 3234–3247. doi: 10.1038/onc.2011.43.

Enzmann, B. L. and Wronski, A. (2018) ‘Synthego’s Engineered Cells Allow Scientists to “Cut Out” CRISPR Optimization [SPONSORED]’, The CRISPR Journal. Mary Ann Liebert, Inc. 140 Huguenot Street, 3rd Floor New Rochelle, NY 10801 USA, 1(4), pp. 255–257.

González-Mariscal, L. et al. (2020) ‘Relationship between apical junction proteins, gene expression and cancer’, Biochimica et Biophysica Acta (BBA)-Biomembranes. Elsevier, 1862(9), p. 183278.

Hirota, C. et al. (2021) ‘Inverse regulation of claudin-2 and -7 expression by p53 and hepatocyte nuclear factor 4α in colonic MCE301 cells’, Tissue barriers. 2020/12/23. United States, 9(1), p. 1860409. doi: 10.1080/21688370.2020.1860409.

Kim, D. and Cho, K.-H. (2023) ‘Hidden patterns of gene expression provide prognostic insight for colorectal cancer’, Cancer Gene Therapy. Nature Publishing Group US New York, 30(1), pp. 11–21.

Kim, H.-Y. et al. (2020) ‘Rumex japonicus Houtt. alleviates dextran sulfate sodium-induced colitis by protecting tight junctions in mice’, Integrative medicine research. 2020/03/03. Netherlands, 9(2), p. 100398. doi: 10.1016/j.imr.2020.02.006.

Li, H., Guo, L. and Cai, Z. (2021) ‘TCN1 is a Potential Prognostic Biomarker and Correlates with Immune Infiltrates in Lung Adenocarcinoma’. Research Square Platform LLC. doi: 10.21203/rs.3.rs-985718/v1.

Mankertz, J. et al. (2004) ‘Functional crosstalk between Wnt signaling and Cdx-related transcriptional activation in the regulation of the claudin-2 promoter activity’, Biochemical and Biophysical Research Communications. Elsevier BV, 314(4), pp. 1001–1007. doi: 10.1016/j.bbrc.2003.12.185.

Mariadason, J. M. et al. (2005) ‘Gene expression profiling of intestinal epithelial cell maturation along the crypt-villus axis’, Gastroenterology. Elsevier BV, 128(4), pp. 1081–1088. doi: 10.1053/j.gastro.2005.01.054.

Meng, H. et al. (2023) ‘Application of CRISPR-Cas9 gene editing technology in basic research, diagnosis and treatment of colon cancer’, Frontiers in Endocrinology. Frontiers Media S.A., 14, p. 1148412. doi: 10.3389/FENDO.2023.1148412/BIBTEX.

Miwa, N. et al. (2001) ‘Involvement of Claudin-1 in the β-Catenin/Tcf Signaling Pathway and its Frequent Upregulation in Human Colorectal Cancers’, Oncology Research Featuring Preclinical and Clinical Cancer Therapeutics. Computers, Materials and Continua (Tech Science Press), 12(11), pp. 469–476. doi: 10.3727/096504001108747477.

Niessen, C. M. (2007) ‘Tight junctions/adherens junctions: basic structure and function’, Journal of investigative dermatology. Elsevier, 127(11), pp. 2525–2532.

Paquet-Fifield, S. et al. (2018) ‘Tight Junction Protein Claudin-2 Promotes Self-Renewal of Human Colorectal Cancer Stem-like Cells’, Cancer Research. American Association for Cancer Research (AACR), 78(11), pp. 2925–2938. doi: 10.1158/0008-5472.can-17-1869.

Roginsky, J. (2018) ‘Analyzing CRISPR Editing Results: Synthego Developed a Tool Called ICE to Be More Efficient Than Other Methods’, Genetic Engineering & Biotechnology News. Mary Ann Liebert, Inc. 140 Huguenot Street, New Rochelle, NY 10801-5215 (914 …, 38(11), pp. S24–S26.

Rosenthal, R. et al. (2010) ‘Claudin-2, a component of the tight junction, forms a paracellular water channel’, Journal of Cell Science. The Company of Biologists, 123(11), pp. 1913–1921. doi: 10.1242/jcs.060665.

Sarkar, S. (2023) ‘Proteogenomic Approaches to Understand Gene Mutations and Protein Structural Alterations in Colon Cancer’, Physiologia. MDPI, 3(1), pp. 11– 29.

Sena, P. et al. (2014) ‘Morphological and quantitative analysis of BCL6 expression in human colorectal carcinogenesis’, Oncology Reports. Spandidos Publications, 31(1), pp. 103–110.

Siegel, R. L. et al. (2023) ‘Colorectal cancer statistics, 2023’, CA: a cancer journal for clinicians. Wiley Online Library, 73(3), pp. 233–254.

Tabariès, S. et al. (2012) ‘Claudin-2 promotes breast cancer liver metastasis by facilitating tumor cell interactions with hepatocytes’, Molecular and cellular biology. 2012/05/29. United States, 32(15), pp. 2979–2991. doi: 10.1128/MCB.00299-12.

Tabariès, S. et al. (2019) ‘Afadin cooperates with Claudin-2 to promote breast cancer metastasis’, Genes & development. 2019/01/28. United States, 33(3–4), pp. 180–193. doi: 10.1101/gad.319194.118.

Tabariès, S. et al. (2021) ‘Claudin-2 promotes colorectal cancer liver metastasis and is a biomarker of the replacement type growth pattern’, Communications biology. Nature Publishing Group UK, 4(1), p. 657. doi: 10.1038/s42003-021-02189-9.

Tao, D. et al. (2023) ‘Expression patterns of claudins in cancer’, Heliyon. England, 9(11), pp. e21338–e21338. doi: 10.1016/j.heliyon.2023.e21338.

Thoma, O. M., Neurath, M. F. and Waldner, M. J. (2021) ‘Cyclin-Dependent Kinase Inhibitors and Their Therapeutic Potential in Colorectal Cancer Treatment. Front. Pharmacol. 12:s757120. doi: 10.3389/fphar.2021.757120’, Frontiers in Pharmacology| https://www.frontiersin.org, 12.

Tian-Yu, Q. et al. (2021) ‘Claudin-14 promotes colorectal cancer progression via the PI3K/AKT/mTOR pathway.’, Neoplasma, 68(5).

Venugopal, S., Anwer, S. and Szászi, K. (2019) ‘Claudin-2: roles beyond permeability functions’, International journal of molecular sciences. MDPI, 20(22), p. 5655.

Wang, D.-W. et al. (2022) ‘The role and mechanism of claudins in cancer’, Frontiers in Oncology. Frontiers, 12, p. 1051497.

Weber, C. R. et al. (2008) ‘Claudin-1 and claudin-2 expression is elevated in inflammatory bowel disease and may contribute to early neoplastic transformation’, Laboratory investigation; a journal of technical methods and pathology. 2008/08/18. United States, 88(10), pp. 1110–1120. doi: 10.1038/labinvest.2008.78.

Wei, C., Xiao-nan, Z. H. U. and Jin, W. (2018) ‘The Expression and Biological Function of claudin-23 in Colorectal Cancer’, Journal of Sichuan University (Medical Science Edition), 49(3).

Wei, M. et al. (2021) ‘Claudin-2 promotes colorectal cancer growth and metastasis by suppressing NDRG1 transcription’, Clinical and translational medicine. United States, 11(12), pp. e667–e667. doi: 10.1002/ctm2.667.

Yang, L. et al. (2022) ‘Evaluation of the Prognostic Relevance of Differential Claudin Gene Expression Highlights Claudin-4 as Being Suppressed by TGFβ1 Inhibitor in Colorectal Cancer’, Frontiers in genetics. Switzerland, 13, p. 783016. doi: 10.3389/fgene.2022.783016.

Yellapa, A. et al. (2014) ‘Interleukin 16 expression changes in association with ovarian malignant transformation’, American Journal of Obstetrics and Gynecology. Elsevier BV, 210(3), pp. 272.e1-272.e10. doi: 10.1016/j.ajog.2013.12.041.

Zhang, X. et al. (2018) ‘CLDN2 inhibits the metastasis of osteosarcoma cells via down-regulating the afadin/ERK signaling pathway’, Cancer cell international. England, 18, p. 160. doi: 10.1186/s12935-018-0662-4.

Zhang, Y. et al. (2015) ‘Tight junction CLDN2 gene is a direct target of the vitamin D receptor’, Scientific reports. England, 5, p. 10642. doi: 10.1038/srep10642.

Zhao, Z. et al. (2021) ‘Review of applications of CRISPR-Cas9 gene-editing technology in cancer research’, Biological Procedures Online 2021 23:1. BioMed Central, 23(1), pp. 1–13. doi: 10.1186/S12575-021-00151-X.

Zhu, L. et al. (2019) ‘Claudin Family Participates in the Pathogenesis of Inflammatory Bowel Diseases and Colitis-Associated Colorectal Cancer’, Frontiers in immunology. Frontiers Media S.A., 10, p. 1441. doi: 10.3389/fimmu.2019.01441.

Zhu, X. et al. (2023) ‘TCN1 Deficiency Inhibits the Malignancy of Colorectal Cancer Cells by Regulating the ITGB4 Pathway’, Gut and liver. 2022/06/10. Korea (South), 17(3), pp. 412–429. doi: 10.5009/gnl210494.

